# Enhanced 5mC-Methylation-Site Recognition in DNA Sequences using Token Classification and a Domain-specific Loss Function

**DOI:** 10.1101/2023.06.01.543218

**Authors:** Wenhuan Zeng, Daniel H. Huson

## Abstract

DNA 5-methylcytosine modification has been widely studied in mammals and plays an important role in epigenetics. Several computational methods exist that attempt to determine the methylation state of a DNA sequence centered at a possible methylation site. Here, we introduce a novel deep-learning framework, MR-DNA, that predicts the methylation state of a single nucleotide located in a gene promoter region. The idea is to adapt the named-entity recognition approach to methylation-site prediction and to incorporate biological rules during model construction. MR-DNA has a stacked model architecture consisting of a pre-trained MuLan-Methyl-DistilBERT language model and a conditional random field algorithm, trained with a self-defined methyl loss function. The resulting fine-tuned model achieves an accuracy of 97.9% on an independent test dataset of samples. An advantage of this formulation of the methylation-site identification task is that it predicts on every nucleotide of a sequence of a given length, unlike previous methods that the predict methylation state of DNA sequences of a short fixed length. For training and testing purposes, we provide a database of DNA sequences containing verified 5mC-methylation sites, calculated from data for eight human cell lines downloaded from the ENCODE database.

## Introduction

DNA methylation is an important epigenetic mechanism. Much attention has been given to 5-methylcytosine (5-mC) modifications in which cytosine is methylated at its fifth position. In mammals, this process is associated with a range of different biological processes, in particular gene regulation, gene silencing, X-chromosome inactivation, genomic imprinting and human cancer development^1–4^.

Several experimental methods have been developed that aim at quantifying 5mC methylation, especially in mammalian genomes, such as bisulfite conversion second-generation sequencing (BS-seq)^5,6^ or using third-generation sequencing technologies, namely Pacbio SMRT sequencing^7^, Nanopore sequencing^8,9^ and Pacbio circular consensus sequencing (CCS), particularly in repetitive genomic regions^10^. Several computational approaches based on machine-learning algorithms have been conceived to detect methylation state on data generated from the above-mentioned sequencing technologies (see Table 1). Most aim at predicting whether the central base of a short DNA sequence is methylated; all show reliable performance^11–14^.

**Table 1.** Overview of work on the 5mC-methylation detection task in the human genome.

| Study | Database | Model architecture | Applied objective |
| --- | --- | --- | --- |
| Wang et al.<sup>17</sup> | Promoter sequences from the UCSC genome browser<sup>18</sup>, iPromoter-5mC database<sup>13</sup>. | Bidirectional Encoder Representations from Transformers (BERT)<sup>19</sup> | DNA fragment of 41 centered on a cytosine |
| Ni et al.<sup>20</sup> | PacBio CCS reads, Illumina and Nanopore data of human sample | Bidirectional Gated Recurrent Unit (GRU) with Bahdanau attention network | Provides both read-level (21-mer sequence that includes CpG in the center) and site-level prediction |
| Wang et al.<sup>21</sup> | iPromoter-5mC database | Transformer encoder and fully connected neural network | DNA fragment of 41 centered on a cytosine |
| Jia et al.<sup>22</sup> | iPromoter-5mC database | A stacked framework of densely connected convolutional networks (DenseNet) and bidirectional GRU with attention mechanism | DNA fragment of 41 centered on a cytosine |
| Jin et al.<sup>16</sup> | Human cell line | Pretrained DNABERT, fine-tuning using adversarial training method | DNA fragments of diverse length centered on a cytosine |
| Bonet et al.<sup>23</sup> | Nanopore data and BS data of Human cell line NA12878 | Convolutional neural network (CNN) | Provides methylation calling on both read-level (length from 1 to 17) and site-level |
| Nyuyen et al.<sup>11</sup> | Human cell line | Machine learning techniques including XGBoost, random forest, deep forest, and deepFNN | DNA fragment of 41 centered on a cytosine |
| Cheng et al.<sup>12</sup> | iPromoter-5mC database | BiLSTM | DNA fragment of 41 centring the cytosine |
| Zhang et al.<sup>13</sup> | RRBS data in cell lines of lung cancer | Deep neural network | DNA fragment of 41 centered on a cytosine |
| Tian et al.<sup>24</sup> | WGBS data of H1 ESC, normal brain white matter, lung and colon tissue. | CNN | DNA fragment of 400 bps centered on a cytosine. |

**Table 2.** Performance evaluation. MR-DNA was trained on three datasets with different sequence lengths. The performance of the methyl loss is compared against that of models trained using a cross-entropy loss function and a focal loss function. For each test dataset, we report accuracy, macro F1-score, macro precision and macro recall. Best values are shown in bold.

| Dataset | Loss function | Accuracy | Macro F1-score | Macro precision | Macro recall |
| --- | --- | --- | --- | --- | --- |
| MR-DNA-50 | Cross-entropy | 0.9780 | 0.8577 | 0.8265 | 0.9206 |
|  | Focal loss | 0.9751 | 0.8557 | 0.8205 | **0.9518** |
|  | Methyl loss | **0.9787** | **0.8659** | **0.8319** | 0.9390 |
| MR-DNA-100 | Cross-entropy | 0.9845 | 0.8323 | 0.7985 | 0.9033 |
|  | Focal loss | 0.9749 | 0.8046 | 0.7708 | **0.9575** |
|  | Methyl loss | **0.9851** | **0.8341** | **0.8012** | 0.8996 |
| MR-DNA-200 | Cross-entropy | **0.9895** | 0.7994 | **0.7699** | 0.8606 |
|  | Focal entropy | 0.9783 | 0.7684 | 0.7394 | **0.9569** |
|  | Methyl loss | 0.9893 | **0.8003** | 0.7695 | 0.8669 |

Previous studies have regarded DNA-methylation identification as a binary classification task. Most of them employ traditional machine learning approaches^11^, classic deep learning techniques such as deep feed-forward neural networks (deepFNN)^11^, bidirectional long short-term memory (BiLSTM) networks^12^, or ensemble frameworks^13,14^ are applied to a combination of features extracted from the DNA sequences. Also, transformer-based language models are trained on DNA sequences to provide a pre-trained model^15^ that is then fine-tuned on a down-stream database for performing DNA-methylation identification^16^. These methods are all designed to predict whether a presented DNA sequence of fixed length 41 or others is methylated in the middle or not. Thus, these methods are not directly applicable to the main practical problem of interest, namely the prediction of individual methylation sites in a sequence.

Hence, there is a need for a machine-learning approach capable of making predictions for individual bases in DNA sequences of arbitrary length. To address this, here we model DNA sequences as “language”, where each 3-mer is a token, and phrase methylation-site prediction as a token classification task, which is a fundamental task in natural language understanding (NLU). The named-entity recognition (NER) task is usually applied in information extraction (IE), for example, to locate characters, events, organizations, or locations^25,26^, or to recognize other types of entities^27,28^ in a text. This approach has been applied to biomedical problems that are focused on text^29–31^, but has not yet been transferred to biological sequences.

Traditional statistical models^32–35^ are gradually being superseded by transformer-based language models or other neural networks^31,36,37^. The stacked model architecture, which allows the stacking of additional layers, such as LSTMs and conditional random fields (CRF), to a pre-trained transformer-based language model, has significantly improved the ability to capture dependencies between entities and corresponding labels^38–40^.

As mentioned, previous approaches address the question of whether the middle base of a DNA sequence of length 41 is methylated or not. In our approach, we take advantage of the fact that NER assigns a label to each token and thus can predict methylation for individual bases in a given sequence. In addition, in order to better adapt general NER approaches to biological scenarios, we propose a novel loss function, methyl loss, that is based on the categorical cross-entropy loss function and uses known biological rules of 5-methylcytosine to help model training.

The MuLan-Methyl framework^41^ consists of five transformer-based language models, each trained on the iDNA-MS database^42^, a comprehensive dataset containing DNA methylation sequences for three methylation types and twelve taxonomic lineages. In this study, we use MuLan-Methyl-DistilBERT, one of five finetuned MuLan models, as the model encoder. We use this particular language model because it outperforms the other constituents of MuLan-Methyl for 5hmC-methylation site recognition in terms of both prediction performance and computational efficiency.

Here, we propose a new model architecture, MR-DNA (<u>M</u>ethylation-site <u>R</u>ecognition in <u>DNA</u>) that consists of MuLan-Methyl-DistilBERT and a conditional random fields (CRF) algorithm. The weights of MR-DNA are updated using methyl loss while training, which enhances the model prediction ability.

To evaluate the performance of MR-DNA, we created a database of DNA sequences, each of length 1000 bp, that correspond to gene promoter regions from eight human cell line projects in the ENCODE database^43,44^, with differing numbers of experimentally-verified methylation sites. We further cut each such DNA sequence into small pieces according to a set stride during model training. The workflow of database construction is shown in Fig. 1A. The performance of the fine-tuned MR-DNA model was evaluated under the exact-match evaluation criterion^26^.

**Figure 1.**
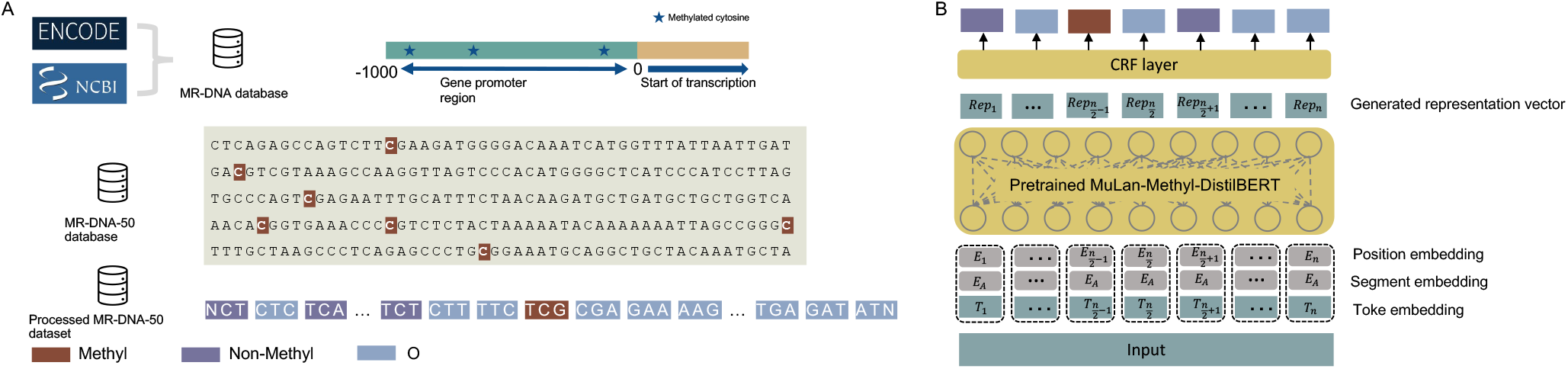
A. Database creation. For eight human cell lines in ENCODE, we extracted 1000 bp upstream gene promoter regions, and annotated cytosines by their reported methylation state. We call this the MR-DNA database. The MR-DNA-50 database was extracted from these sequences using a window size of 50 and stride of 25. The *processed* MR-DNA-50 database was obtained by extracting 3-mers from each sequence in the MR-DNA-50 database and labeling each such 3-mer as methylated, if its central nucleotide is so labeled. B. Model structure. MR-DNA is a stacked model that uses a CRF layer on top of pretrained MuLan-Methyl-DistilBERT, assigning the most probable category to each token.

Our main contributions are:

- This is the first study to phrase methylation-site recognition as a named entity recognition (NER) task.
- We propose a new loss function, methyl loss, that can effectively overcome the impact of skewed data distribution on the model.
- We present a novel framework, MR-DNA, that combines the statistical modelling method CRF with the fine-tuned MuLan-Methyl-DistilBERT model and is trained using methyl loss, showing promising performance on the test dataset.
- We provide a database of DNA gene-promoter sequences (each 1000 bp length) containing varying numbers of verified methylation sites.
- The MR-DNA method aims at recognizing the methylation state of each individual base in a given DNA sequence.

## Results

### Training MR-DNA

MR-DNA is implemented in Python 3.10, using the Pytorch, Huggingface^45^, and pytorch-crf packages. It was run on a Linux Virtual Machine (Ubuntu 20.04 LTS) equipped with 4 GPUs, provided by de.NBI (flavor: de.NBI RTX6000 4 GPU medium). MR-DNA was trained on the above-mentioned processed MR-DNA-50 training and validation datasets using the methyl loss function, configured with a 2e-5 learning rate, early-stopping at 9 epochs, and a batch size of 256 for each GPU.

### Experiment evaluation

Our study uses the exact-match evaluation criterion to assess MR-DNA performance instead of the seqeval^46^ framework designed for sequence labelling evaluation. We use the following standard definitions.

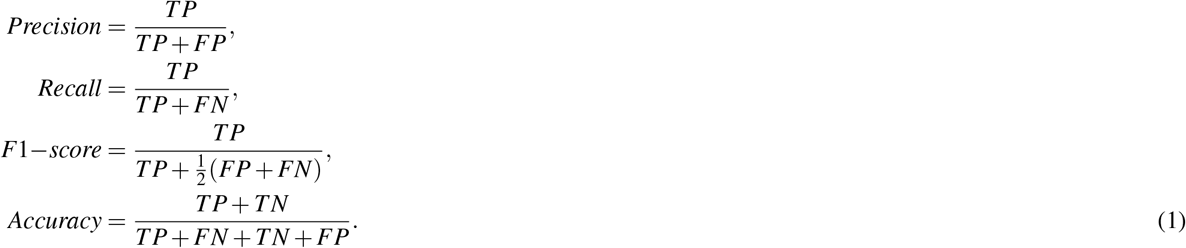

Here, TP, FP, TN, and FN are the true positive, false positive, true negative and false negative counts, respectively. Because the annotations in our database have a highly imbalanced distribution, We use the macro average method to compute the F1-score, precision, and recall, in order to gain a better understanding of the model’s overall performance on imbalanced datasets. This is computed by taking the arithmetic mean of all the class-wise metrics. Each macro-average metric *metric* ∈{*Precision, Recall, F1* − *scores*} is computed as follows:

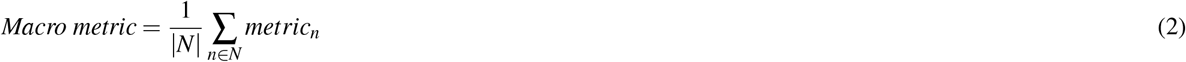

where *N* = {*n*_1_, *n*_2_, …, *n* _*j*_} is the set of classes that appear in the datasets.

### Performance of MR-DNA on multiple datasets with different lengths

First, we investigated the ability of MR-DNA, trained with the methyl loss function, to recognize methylation sites on the processed MR-DNA-50 test data. Here, each sample is a sentence of 50 whitespace-separated entities (3-mers), together with one of three possible annotations (“Methyl”, “Non-Methly” or “O”). We used accuracy, F1-score, recall and precision to quantify the ability of MR-DNA to predict the label of each entity (3-mer). As this is a multiple classification task, we used macro averages to calculate the performance metrics.

Secondly, to explore the effect of sequence length on classification performance, we trained MR-DNA on additional processed MR-DNA-100 and MR-DNA-200 training datasets, as well, using sentences of length 100 and 200, respectively. We observed that MR-DNA with methyl loss trained on MR-DNA-50 outperforms the models trained on the two datasets of longer sequence lengths regarding F1-score, recall, and precision (See Table. 2).

Entities with annotation “O” are easier to identify correctly because they do not have a “C” at the center. To address this, in addition to evaluating model performance when distinguishing between three possible annotations, which are “Methyl”, “Non-Methly” or “O”, we also evaluated model performance distinguishing between only two annotations, “Methyl” and “Non-Methly” (see Table S1).

### Methyl loss enhances MR-DNA performance

We evaluated the usefulness of the proposed loss function on each of the three datasets, MR-DNA-50, MR-DNA-100 and MR-DNA-200, comparing its performance with that of the categorical cross-entropy loss, which we view as a baseline. The datasets statistics of MR-DNA-100 and MR-DNA-200 are shown in Figure S1 and S2.

As shown in Table. 2 and Table S1, for each dataset, MR-DNA trained with the methyl loss outperforms the baseline, especially according to the macro F1-score. Confusion matrices (see Fig. 2) for MR-DNA-50, -100 and -200, show that MR-DNA trained with the methyl loss function on the former outperforms the model trained on the two latter, for the “Methyl” category. Additionally, the model trained with methyl loss has higher sensitivity than the model trained with cross-entropy on the same dataset.

**Figure 2.**
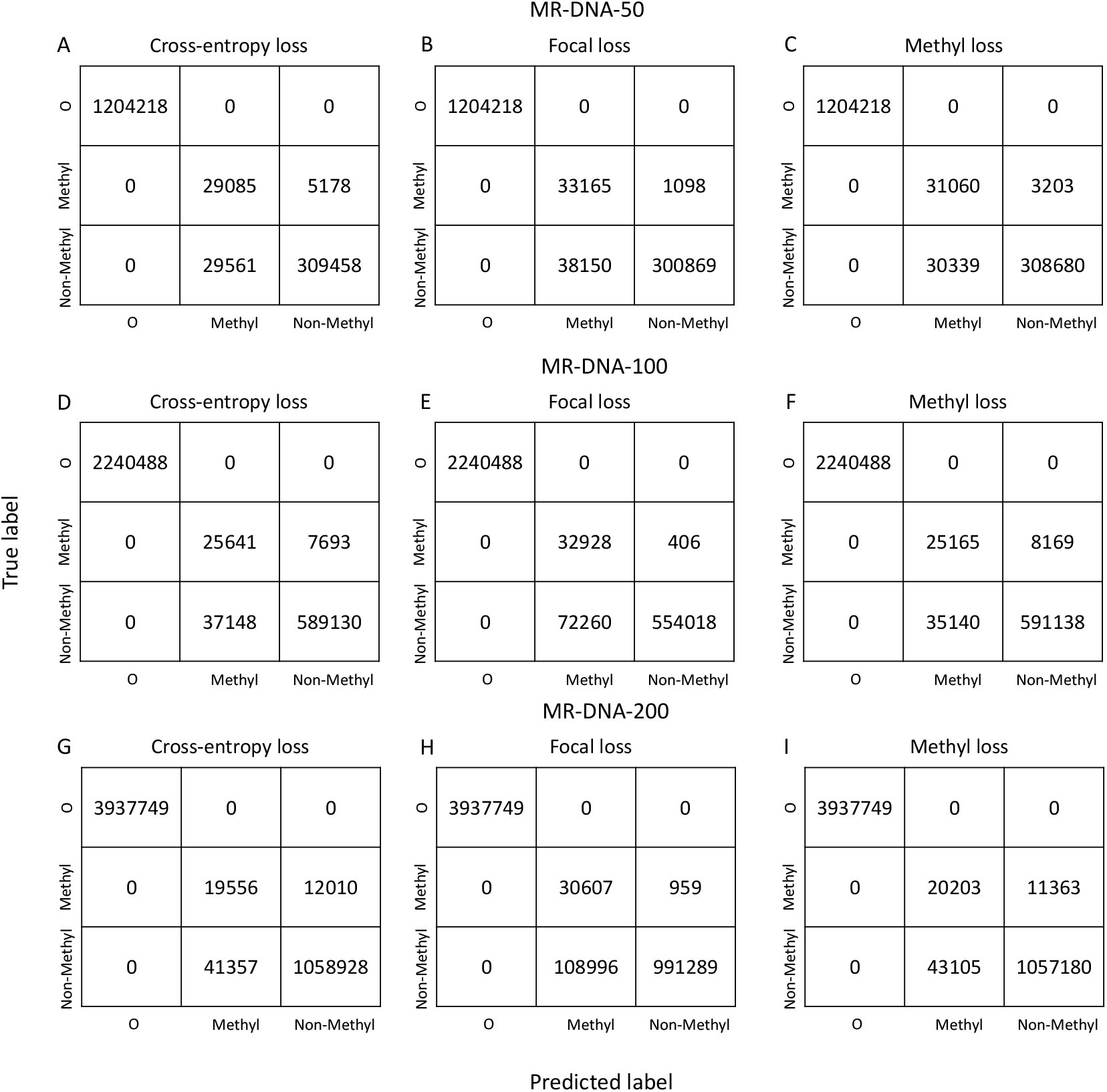
Confusion matrix visualization and comparison of the model trained on (A-C) the MR-DNA-50 training dataset, (D-F) MR-DNA-100 training dataset and (G-I) MR-DNA-200 training dataset using a categorical cross-entropy loss function, a focal loss function, and a methyl loss function, respectively. Performance evaluation was conducted on the corresponding test dataset.

### Methyl loss is less complex and more stable

The focal loss function is widely used in scenarios that train models on an imbalanced dataset. Two hyperparameters, *α* and *γ*, are designed to assign different weights to categories according to their data distribution and control the weight according to the difficulty of predicting the sample, respectively. In comparison, the methyl loss function has one hyperparameter *λ*, used to assign weight according to the commonness of the 3-mer under its label.

We compared the performance of a model trained using the methyl loss function and one trained using the focal loss function on each of the three datasets. The *α* and *γ* values for focal loss are [0.1, 0.7, 0.2] and 5, respectively.

For all three datasets, the model with focal loss function performs worse than the one with methyl loss with respect to accuracy, macro f1-score, and macro precision (see Table. 2). Using the hyperparameter *α*, the focal loss function puts more attention on the least category ‘Methyl’, however, it also leads to a poor false positive rate for this category as shown in Fig. 2. In comparison, the methyl loss function is more stable, improving the model performance in all categories.

Notably, as we mentioned above, we computed evaluation metrics in two ways, one is obtained by considering models’ overall performance in identifying three annotations of entities (See Table. 2 and Table. S1), and another is only considering models’ performance in identifying “Methyl” and “Non-Methly” entities (Table. S1). Comparing these two tables, the values in Table. S1 are comprehensively lower than the value in Table. 2, which is caused by excluding the promising performance in identifying “O” entities. however, the relative differences between the performance of the different models are consistent.

### Stability and flexibility of MR-DNA

Our study is the first one that aims to predict the methylation state of every base in a given DNA sequence. Its novelty makes it difficult to compare to previous studies, which aim at predicting the global methylation state of DNA sequences with fixed length, where the target cytosine lies in the center of the sequences. To overcome this, we modified the prediction format output by MR-DNA to allows comparisons against previous studies using the iPromoter-5mC database^13^ as the benchmark database, and applying our models to the independent test dataset of the benchmark database, namely the iPromoter-5mC test dataset.

The iPromoter-5mC test dataset consists of DNA sequences with a length of 41, labelled ‘positive’ if the centerd cytosine is methylated, and ‘negative’, otherwise. The DNA sequences in the iPromoter-5mC test dataset lack methylation state annotations for all bases except for the one located in the center.

To allow model comparison, the iPromoter-5mC test dataset was processed by converting the DNA sequence to a sequence that consists of 41 consecutive 3-mers, using the same processing approaches as the one used for our database. We applied the MR-DNA-50, -100, -200 that were trained on the corresponding MR-DNA database directly to the processed iPromoter-5mC test dataset for predicting each DNA sequence’s global methylation state, respectively. MR-DNA is not trained on the training dataset of the iPromoter-5mC database since its data annotation is inadequate for training the MR-DNA model. Additional, we converted the output format of the NER task to the output format of the binary classification task, in which each DNA sequence is assigned ‘1’ if the central 3-meris predicted as ‘Methyl’, and ‘0’, otherwise.

We calculated accuracy, f1-score and specificity to evaluate our models’ performance on the iPromoter-5mC test dataset, and for comparing against models from previous studies which are trained and tested on the iPromoter-5mC training dataset. Some evaluation metrics were computed using the weighted-average approach due to the imbalanced distribution of the iPromoter-5mC test dataset.

As shown in Fig. 3, among the MR-DNA models, the MR-DNA-100 performs best regarding recall, specificity, and accuracy. However, in comparison to the best performance model in previous studies, its accuracy, specificity, and recall are 10%, 5.4%, and 2.4% lower, respectively. We emphasise that the models we compared against were trained on the training set that corresponds to the iPromoter-5mC test dataset, while the MR-DNA models were trained on our own dataset, which was collected from different resources than the iPromoter-5mC test dataset. So, despite the difference between our training dataset and iPromoter-5mC test dataset, our models, especially MR-DNA-100, shown comparable performance on the iPromoter-5mC test dataset.

**Figure 3.**
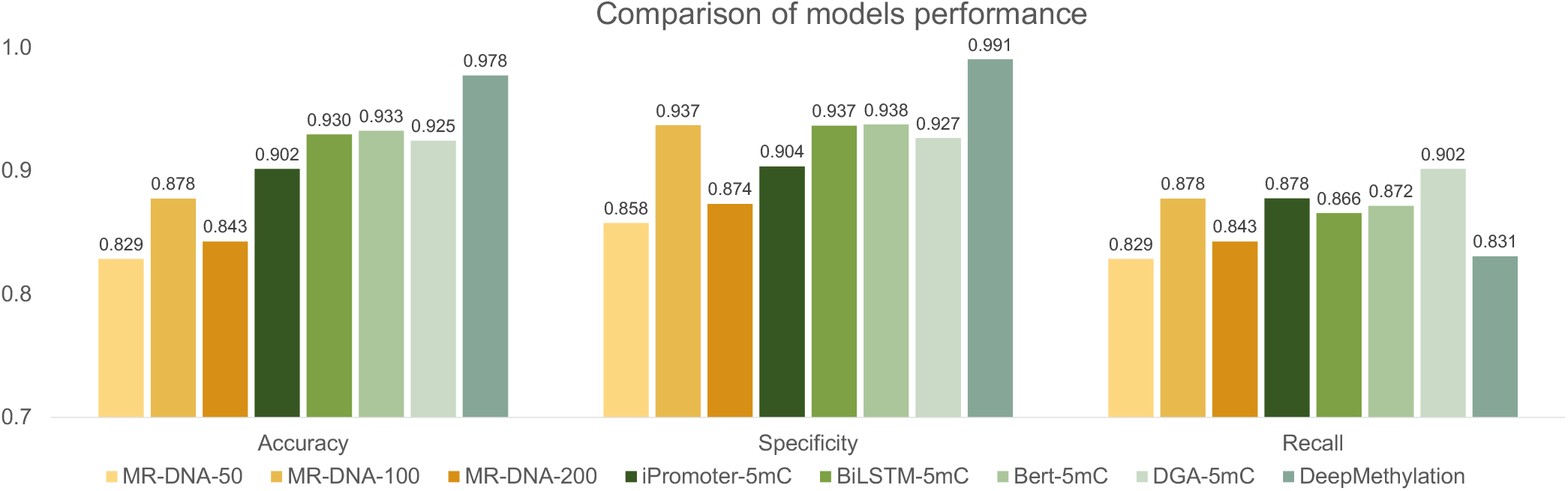
Comparison of model performance between MR-DNA and previous studies. The performance comparison is conducted by comparing the evaluation metrics computed by our models, namely MR-DNA-50, -100, -200, and those computed by previous studies, namely iPromoter-5mC and Bert-5mC, on the iPromoter-5mC test dataset.

## Discussion

The study of 5mC-modifications in mammalian genomes helps the understanding of several biological processes, such as gene expression control. Previous computational approaches based on machine-learning algorithms for identifying methylation sites are only able to categorize DNA sequences of a predetermined length rather than detect methylation sites of a single nucleotide, resulting in limited practical use.

Here, we presented MR-DNA, a novel framework based on natural language understanding technology, which transforms a naive classification task for DNA sequences into a NER problem with the ability to identify the methylation state of each entity. We enhanced MR-DNA’s classification ability by using a custom loss function, methyl loss, to update model weights during training. The sensitivity of classifying the minority category is improved by optimizing the loss function, so as to put more attention on challenging data by adding an exponential based on the categorical cross-entropy loss function. This work shows how machine-learning approaches may be used to determine the methylation state of a specific cytosine, and allows the application for methylation site identification on longer DNA sequences.

The evaluation of MR-DNA’s performance and the effectiveness of methyl loss was conducted using four different metrics compared with the model using the default loss function. Methyl loss shows superior performance in most comparisons among models trained on different datasets. Using methyl loss, all four evaluation metrics indicate that MR-DNA-50,in particular, shows high reliability. We illustrated how one might combine a pretrained language model and a classic statistical algorithm to improve predictive ability.

Moreover, by modifying the output of MR-DNA models, we were able to estimate how well our models work compared to other studies that focus on predicting the methylation state of a given DNA sequence, where the target cytosine is located in the center. The comparison demonstrated that our models, especially the MR-DNA-100 model have comparable prediction ability with the models that are specifically trained for predicting the global methylation state of DNA sequence. So, while the main aim of MR-DNA is to determine the methylation state of individual nucleotides, it also performs well when used to predict the global methylation state of a DNA sequence.

MR-DNA is the first study to transfer the NER approach from human language to biological sequences and to adapt the recognition target to the DNA methylation scenario, aiming to recognize the entities of “methylation”, “non-methylation” and “other nucleotide” in DNA sequences, instead of “persons”, “locations”, and “organizations”, in text.

Additionally, this study offers a custom MR-DNA database generated by obtaining methylation state from eight human cell lines in the ENCODE database, each record consists of 1000 bp in the original MR-DNA database. MR-DNA was trained on the processed dataset by extracting different sequence lengths following self-define standards. The MR-DNA database is provided as a benchmark dataset for researchers interested in optimizing methylation-site detection.

We would like to emphasize that, for a sequence of arbitrary length *n*, a prediction is made by partitioning the sequence into *n* − 50 + 1 individual 50-mers and then applying MR-DNA to each such sequence.

## Methods

### Database construction

The MR-DNA database was constructed from DNA methylation-site information of eight human cell lines (K562, HepG2, GM12878, HeLa-S3, A549, SK-N-SH, H1, GM23248) downloaded from the ENCODE portal^43,44^, with the following identifiers: ENCSR765JPC, ENCSR786DCL, ENCSR890UQO, ENCSR550RTN, ENCSR481JIW, ENCSR145HNT, ENCSR617FKV, ENCSR625HZA. Each experiment provided the methylation state at CpG, CHH, CHG locations, determined using whole-genome bisulfite sequencing (WGBS), in BED format. We filtered the BED files to keep only methylation sites that are on the same strand as the gene, have sequencing coverage of 10 or more, and for which 100% of the assembled reads are reported as methylated. For each of the human cell lines, the filtered BED files were split in a ratio of 9:1 to generate training and test sets, respectively.

The start and end positions of gene promoter regions were determined from the GRCH38 annotation file downloaded from NCBI as the 1000 bp upstream of a transcription start position, and the corresponding DNA sequences were extracted from the GRCH38 genome reference. The filtered BED files were used to annotate the methylated sites in the promoter regions. The methylation site statistics on annotated gene promoter regions in terms of each human cell line project are reported in Fig. 4A-B and Table. S2.

**Figure 4.**
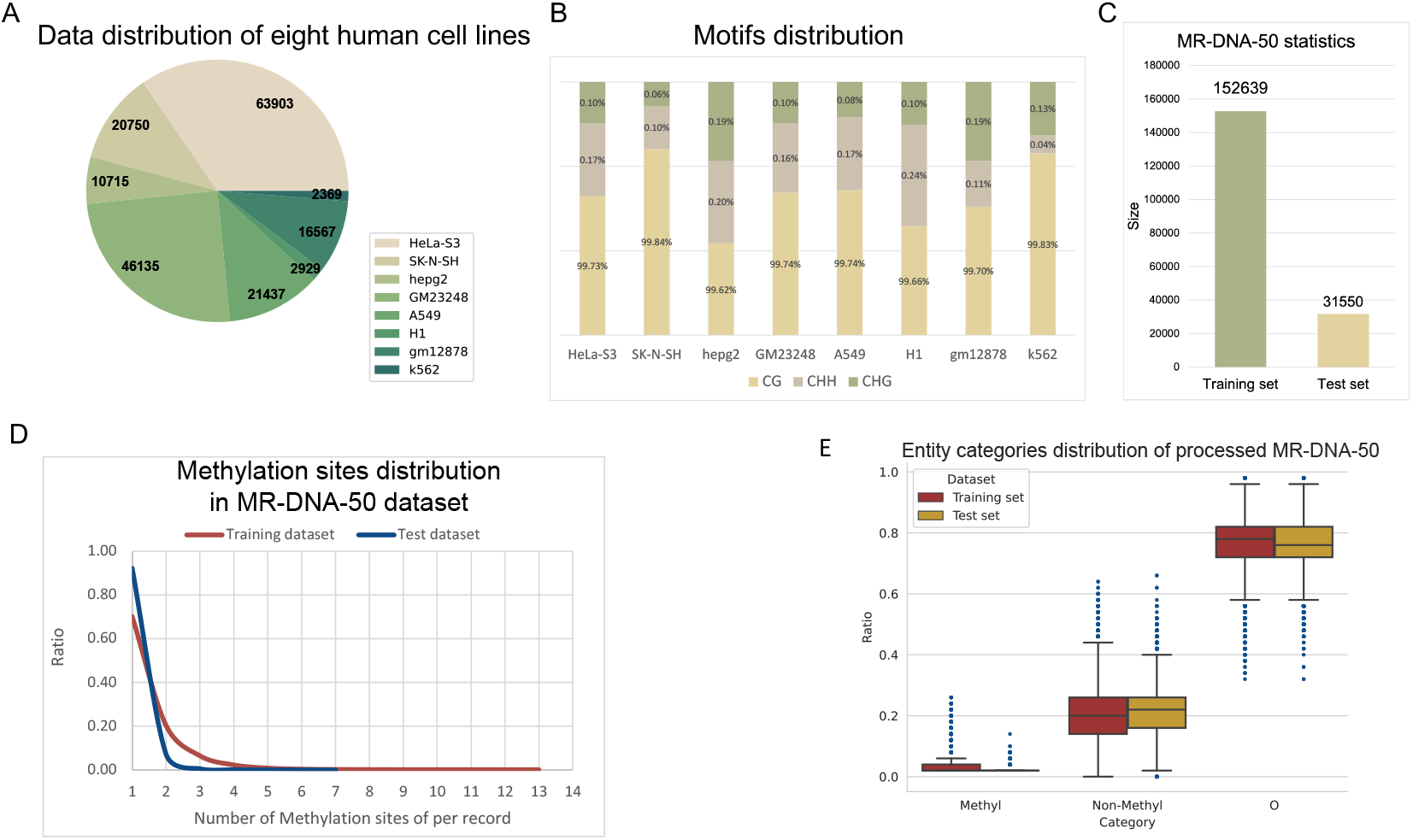
Database statistics. For annotated gene promoter regions, we show (A) a pie chart of the portion of each human cell line project in the annotated gene promoter regions, (B) the distribution of three types of methylation motifs of each human cell line project. For both the training and test dataset of MR-DNA-50, we show (C) a comparison of their sizes, (D) the distribution of ratios of methylation sites per sample, and (E) a box-plot of the entity category distributions.

We first populated the MR-DNA database with the described annotated promoter sequences of length 1000 bp. Then, for training purposes, we constructed a second database, MR-DNA-50, in which all sequences were cut into short sequences of length 50 bp (50-mers), using a stride of 25. Any such 50-mer that is annotated with at least one methylation site was used to generate the MR-DNA-50 training dataset, and the MR-DNA-50 test dataset, respectively (see Fig. 4C).

### Entity category annotation

In preparation of the NER task, we processed and annotated all 50-mers as follows. First, each 50-mer was extended in either direction by a single “N” and each resulting 52-mer was converted into a sentence-like sequence consisting of 50 (=52-3+1) overlapping 3-mers obtained using a sliding window, separated by whitespace. Second, imitating the named-entity naming rule, for each 3-mer, we attached the label “Methyl”, ‘Non-Methyl”, or “O”, depending on whether the middle base is a methylated cytosine, a non-methylated cytosine, or neither, respectively. It is worth mentioning that every nucleotide will appear in the center of some 3-mer, so while a methylated cytosine might be regarded as Non-Methyl when encountered off-center, it will be annotated as Methyl when encountered in the center of a 3-mer.

In result, in both the processed MR-DNA-50 training dataset and test dataset, each sample is a sentence consisting of 50 individual 3-mers separated by whitespace, where each element is tagged as “Methyl”, “Non-Methyl”, or “O”, depending on its methylation state reported in the MR-DNA database (see Fig. 1A).

The median value of the number of methylation sites for each sample in the initial MR-DNA-50 training dataset is 1; thus the number of “Methyl” entities is smaller than the number of “Non-Methyl” entities, which in turn is much smaller than the number of “O” entities (see Fig. 4D-E), and so the distribution of labels is skewed.

### Pre-trained MuLan-Methyl-DistilBERT

Our proposed framework, MR-DNA, consists of a token-level classifier based on the pre-trained MuLan-Methyl-DistilBERT^41^, followed by a Linear-Chain CRF as its decoder layer (see Fig. 1B).

The MuLan-Methyl-DistilBERT classifier is an adaption of DistilBERT^47^ to the problem of DNA-methylation identification, obtained by pre-training the masked language modelling (MLM) task on a custom corpus consisting of both DNA methylation and taxonomy data, and then fine-tuning on a DNA methylation-site dataset. The motivation here is that sequence representation vectors encoded by MuLan-Methyl-DistilBERT that were trained taking domain adaption in to account can better capture the potential semantic relationships and information between tokens.

MuLan-Methyl-DistilBERT uses the same basic neural network architecture as DistilBERT, performing knowledge distillation of Bidirectional Encoder Representations from Transformers (BERT)^19^, while reducing the number of transformer layers and adjusting pre-training tasks. The training objective of DistilBERT is a linear combination of distillation loss ℒ_*ce*_, language modelling mask loss *ℒ*_*mlm*_, and cosine-distance loss *ℒ*_*cos*_^47^ as (3):

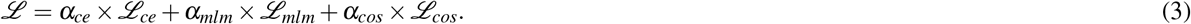

We now describe how to obtain representation vectors. Note that the pre-trained MuLan-Methyl-DistilBERT was trained on a custom corpus and that the vocabulary of the corresponding custom tokenizer contains all combinations of A, T, G, C of multiple lengths as well as words related to taxonomy lineages. However, the DNA sequences in the processed MR-DNA-50 dataset contain an additional character, N. So, the first step is to expand the vocabulary size of the original tokenizer from 25,000 to 25,032, where the additional 32 words are the combinations of A, T, G, C, N of length 3 that occur in the processed MR-DNA-50 training dataset. The embedding dimension of the pre-trained language model was expanded accordingly.

After tokenizing the MR-DNA-50 training dataset using the expanded tokenizer, we employed the updated pre-trained MuLan-Methyl-DistilBERT as an encoder to obtain an embedding representation vector in a three-dimensional array *W* ∈ ℝ^1×*t*×768^, for each input sequence *X* = {*x*_1_, *x*_2_, …, *x*_*t*_}, where *t* is the number of tokens.

### Linear-chain CRF layer

For each input sequence, the embedding representation vectors *W* (generated by the encoder) serve as the input for the linear-chain CRF algorithm, for which the posterior probability *y* of the input sequence is defined as (4):

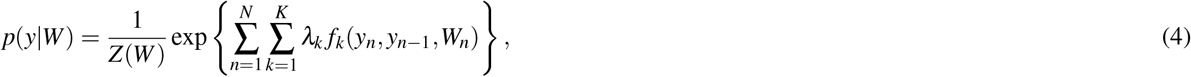

where 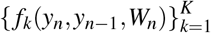 is a set of real-valued feature functions, {*λ*_*k*_} ∈ ℝ^*K*^ is a parameter vector, and *Z*(*W*) is a normalization factor of the form

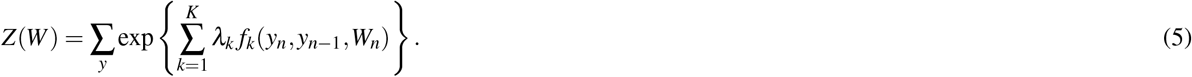

The objective for parameter learning is to maximize the conditional likelihood of the training data given by

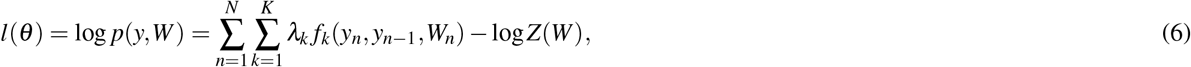

and the most probable assignment for each element in the input sequence

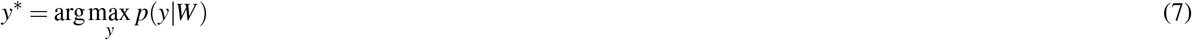

is further decoded by the Viterbi algorithm.

### Methyl loss function

Because of the skewed distribution of entity categories, the model’s ability to predict minority entities is comparatively weak. To mitigate this, we propose to use “methyl loss”, a loss function that allows the model to focus more on challenging samples by embedding the biological rule of DNA methylation.

Methyl loss is based on the categorical cross-entropy loss function,

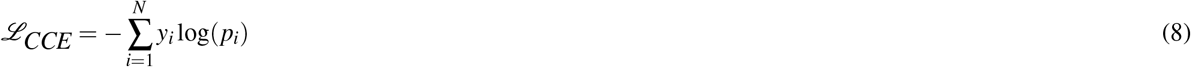

where *y*_*i*_ is the truth label, *p*_*i*_ is the predicted probability of the *i*_*th*_ class. This is inspired by focal loss, which is a variant of cross-entropy loss that has been widely used in the context of imbalanced data distribution:

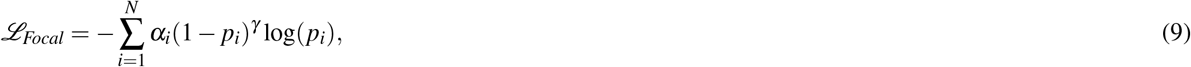

where *γ >* 0 is used to reduce the relative loss for entities well-classified during model training and to concentrate on challenging entity categories. Here, *α*_*t*_ is a hard hyper-parameter containing a list of weights corresponding to each category. The assignment of *α* and *γ* only depends on the ratio of classes, regardless of the scenario.

Methyl loss closes this gap by utilizing the generated annotation vector, for each sample *X* = {*x*_*t*_}_*t*∈*T*_, where *T* is the number of tokens, and *x*_*t*_ is a token consisting of three elements. We define the annotation vector *A* = {*a*_*t*_}_*t*∈*T*_ as

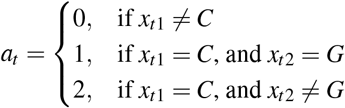

We assume a token is difficult to classify if its nucleotides belong to the minority situation of its category. For instance, each token contains three nucleotides, whereas most tokens with the “Methyl” label contain CpG, where C is in the center. The proposed methyl loss aims to pay more attention to the token which belongs to the minority of its category during model training. In consequence, the methyl loss function is given by

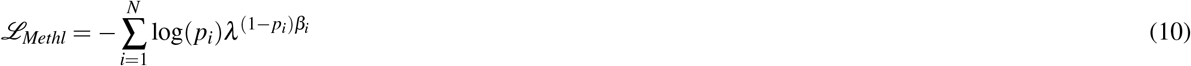

where

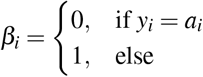

and *λ >* 0 is a variable for controlling the attention level on challenging objects. For choosing the optimal *λ*, we evaluate the impact of *λ* on model performance by setting *λ* from 2 to 5, where *λ* equals 2, resulting in the best accuracy and F1-score in our scenario.

## Data availability

The data used in this study were downloaded from the ENCODE database at https://www.encodeproject.org/. The derived MR-DNA datasets and our code are available at https://github.com/husonlab/MR-DNA.

## Acknowledgements

We acknowledge the support of the BMBF-funded de.NBI Cloud within the German Network for Bioinformatics Infrastructure (de.NBI) (031A532B, 031A533A, 031A533B, 031A534A, 031A535A, 031A537A, 031A537B, 031A537C, 031A537D, 031A538A).

## Author contributions statement

W.Z. conceived and conducted the experiments, W.Z analysed the results. W.Z. and D.H.H. wrote and reviewed the manuscript.

## Additional information

The corresponding author is responsible for submitting a competing interests statement on behalf of all authors of the paper.

